# The diversity of antibiotic resistance and virulence genes are correlated in human gut and environmental microbiomes

**DOI:** 10.1101/298190

**Authors:** Pedro Escudeiro, Joël Pothier, Francisco Dionisio, Teresa Nogueira

**Affiliations:** cE3c – Centro de Ecologia, Evolução e Alterações Ambientais, Faculdade de Ciências, Universidade de Lisboa, Lisbon, Portugal; Atelier de Bioinformatique, ISYEB, UMR 7205 CNRS MNHN UPMC EPHE, Muséum National d’Histoire Naturelle, Paris, France

## Abstract

Human beings have used large amounts of antibiotics, not only in medical contexts but also, for example, as growth factors in agriculture and livestock, resulting in the contamination of the environment. Even when pathogenic bacteria are the targets of antibiotics, hundreds of non-pathogenic bacterial species are affected as well. Therefore, both pathogenic and non-pathogenic bacteria have gradually become resistant to antibiotics. We tested whether there is still co-occurrence of resistance and virulence determinants. We performed a comparative study of environmental and human gut metagenomes issuing from different individuals and from distinct human populations across the world. We found a great diversity of antibiotic resistance determinants (ARd) and virulence factors (VFd) in metagenomes. Importantly there is a correlation between ARd and VFd, even after correcting for protein family richness. In the human gut there are less ARd and VFd than in more diversified environments, and yet correlations between the ARd and VFd are stronger. They can vary from very high in Malawi, where antibiotic consumption is unattended, to non-existent in the uncontacted Amerindians. We conclude that there is co-occurrence of resistance and virulence determinants, suggesting a possible co-selective mechanism. For example, by selecting for resistant bacteria, we may be selecting for more virulent strains as a side effect of antimicrobial therapy.

## Importance

Every year thousands of tons of antibiotics are used, not only in human and animal health but also as growth promoters in livestock. Consequently, during the last 75 years, antibiotic resistant bacterial strains have been selected in human and environmental microbial communities. This implies that, even when pathogenic bacteria are the targets of antibiotics, hundreds of non-pathogenic bacterial species are also affected. Here, we performed a comparative study of environmental and human gut microbial communities issuing from different individuals and from distinct human populations across the world. We found that antibiotic resistance and pathogenicity are correlated and anticipate that, by selecting for resistant bacteria, we may be selecting for more virulent strains, as a side effect of antimicrobial therapy.

## Introduction

Antibiotics are present in microbial communities, not only as a result of the natural lifestyle of microorganisms but also due to the usage of these drugs in agriculture, food industry, livestock, or in human health (1). Therefore, antibiotics can affect bacterial communities as a whole, comprising both pathogenic and non-pathogenic bacteria. Take, for example, the human microbiome, defined as the set of microorganisms that colonize humans. This microbiome is composed of about 3.8e+13 bacterial cells (2) spanning thousands of taxa and colonizing our body’s surfaces and bio fluids, including tissues such as skin, mucosa and most importantly, the gastrointestinal tract. Thus, even when virulent bacteria are the targets of antibiotics, the administration of these drugs may also affect many non-pathogenic mutualistic or commensal bacterial species present in individuals undergoing treatment (3).

The resistome (the collection of all antibiotic resistance (AR) genes, which exist in both pathogenic and non-pathogenic bacteria (4) is frequently encoded on mobile genetic elements. Similarly, the virulome, the set of genes encoding virulence can also be encoded on the mobilome (5–7). Therefore, many bacterial virulence factors (VF) are easily spread in bacterial populations by horizontal gene transfer, converting mutualistic or commensal bacteria into potential opportunistic pathogens. Several examples of highly virulent and multi-resistant disseminated clones have already been reported throughout the literature (for a review see (8)).

A given bacterial population may constitute a life insurance to other bacterial populations present in the same microbiome through at least two different mechanisms. First, if antibiotic resistance genes are coded by plasmids or other mobile genetic elements, they may transfer to pathogenic cells and save them from the negative effects of antibiotics. Indeed, these genetic elements can spread into the bacterial community by horizontal gene transfer, even crossing species (9–11). Curiously, some bacterial strains from different species have been shown to be extremely good donors of certain plasmids. As such, they are able to amplify the number of plasmids in a bacterial community and spread those plasmids to other bacterial cells (12), a phenomenon probably explained by interactions between different plasmids (13, 14).

Secondly, even bacterial cells not coding for antibiotic resistance determinants nor receiving mobile elements may be protected by cells coding for certain drug resistance genes. Indeed, gene products that inactivate antibiotics by degrading or modifying antibiotic molecules are also decreasing their concentration in the local environment (for a review about mechanisms of possible indirect resistance see (15)). This mechanism of indirect resistance has been shown to occur in different systems and to be pervasive (16–20).

In this paper we ask whether the abundance of antibiotic resistance protein families correlate with that of virulence, among bacterial populations. For this, we have chosen to study metagenomes. There are three main reasons to use metagenomes to address this question. First, it is known that, for millions of years, bacteria had to cope with the presence of many other species, mostly competing with them (21), but also cooperating, both phenomena relying on the full set of genes of the metagenome (21). Secondly, horizontal gene transfer promotes the genetic relationship between species thus enforcing cooperation (6) avoiding the emergence of cheaters within the microbiome. Third, the study of metagenomes gives us access to the repertoire of genes involved in adaptation to the environment, given that many of these traits are often encoded in the mobilome and thus can be shared by different, eventually unrelated bacteria. In this context, mining for genes coding for antibiotic resistance and virulence traits in metagenomes is a reliable way to access the selective pressures the population is subject to, as well as the co-occurrence of genetic traits of the whole microbiome.

The present study aims at understanding the relationship between antibiotic resistance genes and those coding for virulence. Here we show that there is indeed a linkage between the dissemination of virulence factors and genes coding for antibiotic resistance, within natural microbiomes, and that this relationship could be influenced by the behaviours of human populations, spanning from very different geographical locations across the world.

## Results

### Antibiotic resistance (AR) gene families in the metagenomes

We started by evaluating the quantity of different AR proteins present in the metagenomes under study. We wanted to know how does the diversity of antibiotic resistance gene families’ homologues (that is, ARd, the number of different protein families) vary with the metagenome protein family richness (that is the total number of proteins families in a metagenome). To answer this question, we used a data set of environmental metagenomes issuing from diverse ecosystems and biomes, such as oceans, coral atolls, deep oceans, Antarctic aquatic environments and snow, soils, hyper saline sediments, sludge’s, microbial fuel cell biofilms, and animal microbial populations (22). In this dataset, there is a broad variation in AR gene diversity (ARd) (Fig 1A). However, there is a correlation between ARd and the protein family richness of the various metagenomes (overall correlation of 0.754). In Fig 1A, regression lines for human gut microbiome (in blue, slope 0.0037) and for other biomes (in black, slope 0.0052) are shown. Nevertheless, the p-value for the difference of these slopes is 1.4e-02, which is significant but not highly significant, does not allow, in our opinion, to draw solid conclusions from this difference.

**Figure 1.**
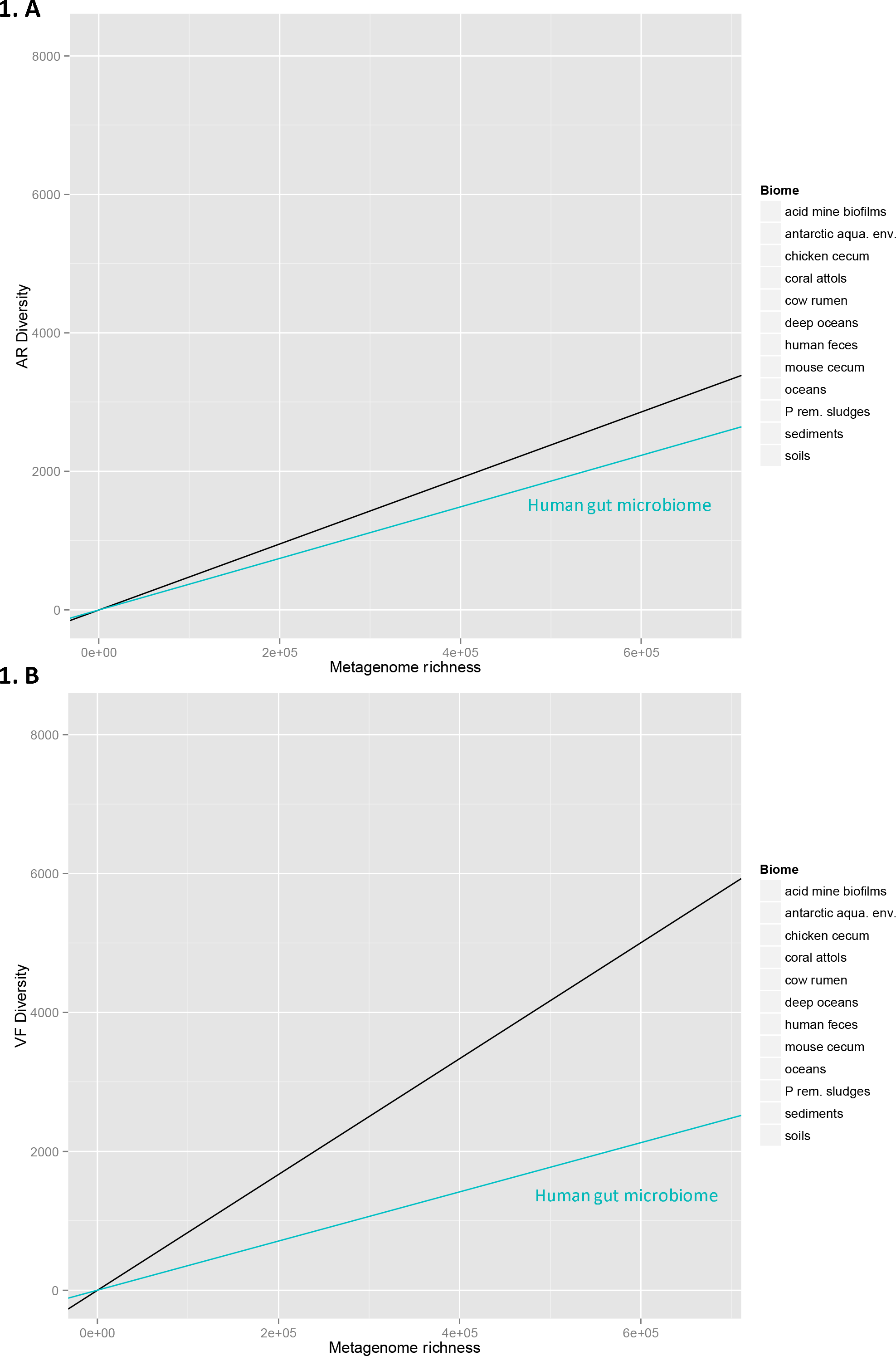
Distribution of the diversity number of antibiotic resistance and virulence factor protein families by environmental metagenomes’ protein family richness. The vertical axes represent the total diversity count of AR protein families’ homologues (ARd) (1. A) and VF protein families’ homologues (VFd) (1.B), present in metagenomes. The horizontal axes represent the protein family richness of the metagenome, that is, the number of cluster representative sequences - see Methods. Each dot represents one of the 64 metagenomes. In 1.A (resp.1B), the black lines represent the regressions of the ARd (resp. VFd) of all the metagenomes on the metagenome richness. The points are scattered showing that the diversity of AR and even more the diversity of VF gene families can vary greatly from metagenome to metagenome. The light green lines represent the linear regressions of the ARd (resp. VFd) of the human faeces metagenomes subset on metagenome richness.

**Figure 2.**
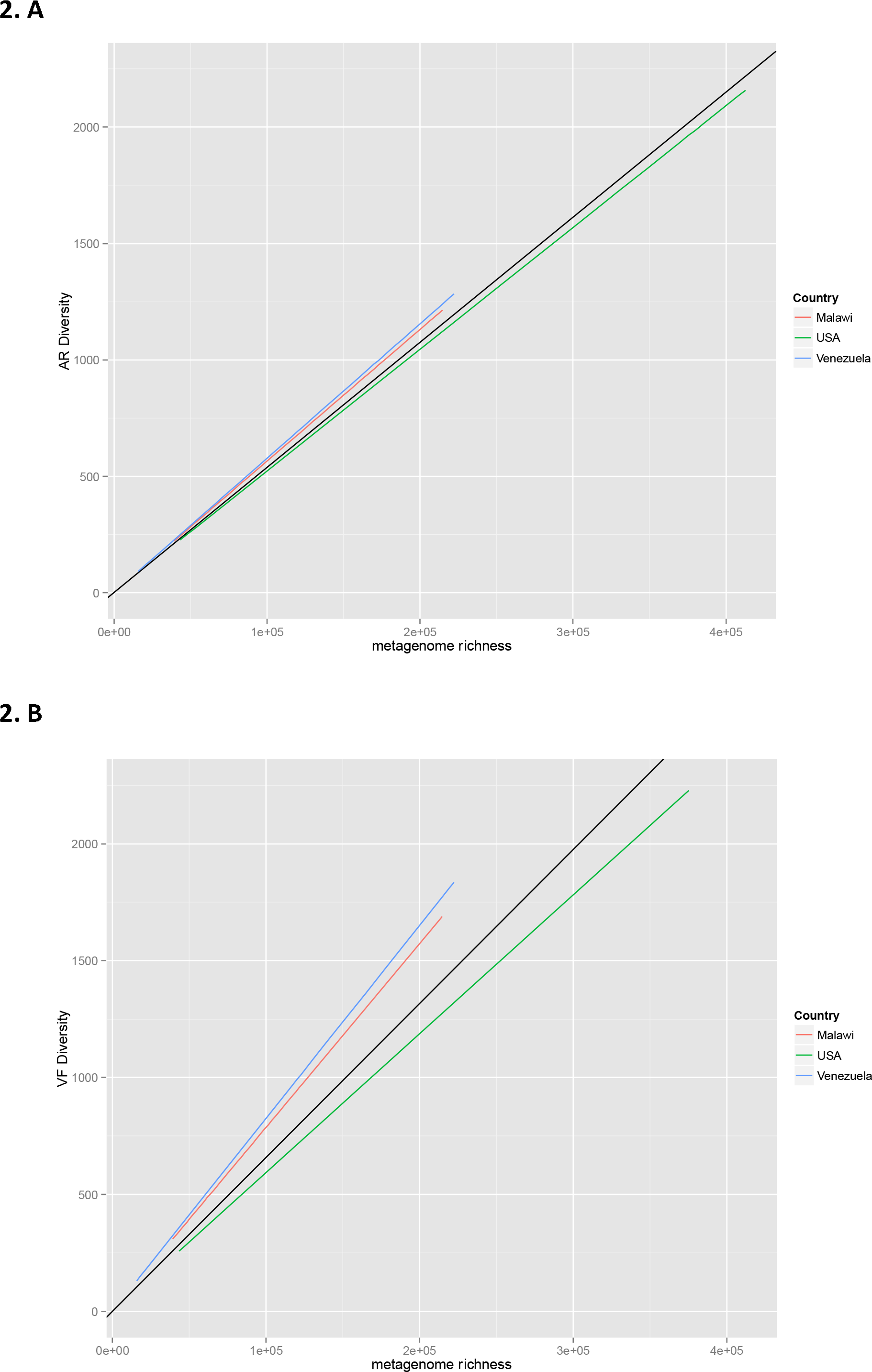
Distribution of the diversity number of antibiotic resistance and virulence factor protein families in human gut metagenomes versus metagenome protein family richness. The vertical axes represent the total diversity count of AR protein families’ homologues (ARd) (2. A) and VF protein families’ homologues (VFd) (2.B), present in the 110 human gut metagenomes (see Material and Method). As in Figure 1, the horizontal axes represent the protein family richness of the metagenomes. Each dot represents one of the 110 human gut metagenomes. In 2.A (resp.2B), the black lines represent the regressions of the ARd (resp. VFd) of all the metagenomes on the metagenome richness. The green lines represent the linear regression of the ARd (resp. VFd) of the USA metagenomes subset on metagenome richness, the red lines are regression lines for the Malawian metagenomes, and the blue lines are those for the Venezuelan metagenomes.

Another dataset composed by 110 human gut metagenomes, sampled from healthy individuals with ages ranging from 0.05 to 53 years of age, spanning different regions of the world such as: US, Malawi, and the Venezuelan Amazon was also studied (23).

### Virulence factor (VF) gene families in the metagenomes

Bacterial virulence factors are proteins that enable pathogenic bacteria to parasitize a host, including gene products involved in adhesion and invasion, secretion, toxins, and iron acquisition systems (24).

The environmental microbiomes of our dataset reveal a great diversity of virulence factors protein families’ (VFd) densities (Fig 1B). Since virulence may also be associated with the colonization of different types of biomes (some virulent traits are involved in adaptation to new environments, by allowing bacteria to adhere and colonize substrates, to access to resources such as iron, among other functions) besides the context of infection, one can expect different types of these genes in environmental microbiomes. Moreover, Fig 1B shows that in human gut microbiomes, VFd versus protein family richness of the metagenomes achieves an overall correlation coefficient of 0.669. Regression lines for human gut microbiome (in blue, slope 0.0035) and for other biomes (black, slope 0.0106) are shown. Here the p-value for difference of these slopes is 1.7e-08, indicating that there could be a difference, but the points are very spread over all types of biome, and we prefer not to draw conclusion about these data.

### The AR / VF correlations

The main purpose of the present work is to evaluate the relationship, if any, between ARd and VFd in microbiomes. Therefore, we excluded from our analysis all the gene products that are both homologues to AR and VF determinants. Thus, we avoid the introduction of a bias due to both correlations of AR and VF diversities (ARd and VFd) with metagenomes richness. In the analysis, we systematically analysed the residuals of the regression of ARd on metagenome richness and the residuals of the regression of VFd on metagenome richness. In Fig 3 one can see that the residuals of ARd and those of VFd are positively correlated. Although there is a large variation of ARd (Fig 1A), and VFd (Fig 1B) in environmental metagenomes, there is a strong correlation between ARd and VFd (Fig 3): the overall partial correlation between ARd and VFd given metagenome richness is 0.84 (p-value 0.8e-35~i.e. 0.0). Indeed, the partial correlation between ARd and VFd given the metagenome richness is the same as the correlation between residuals of ARd and residuals of VFd (the partial correlation coefficient for environmental biomes is 0.86 while the one for the human gut is 0.90). When we draw the regression line for the human gut microbiomes from this dataset (human gut), we can see that the ratio of (residuals ARd)/(residuals VFd) is higher with slope of 0.63 than for the case of environmental metagenomes with slope of 0.44. The difference in slopes is significant, but not very significant, with p-value of 3e-02. Despite the fact that the values for ARd and VFd are lower in gut metagenomes (see Fig 1), one can witness a higher ratio ARd/VFd (larger slope) of metagenomes pertaining to the human faeces biome relatively to the slope calculated for all environmental metagenomes (Fig 3). This suggests that there could be a greater accumulation of ARd over VFd for this particular biome than in environmental ones.

**Figure 3.**
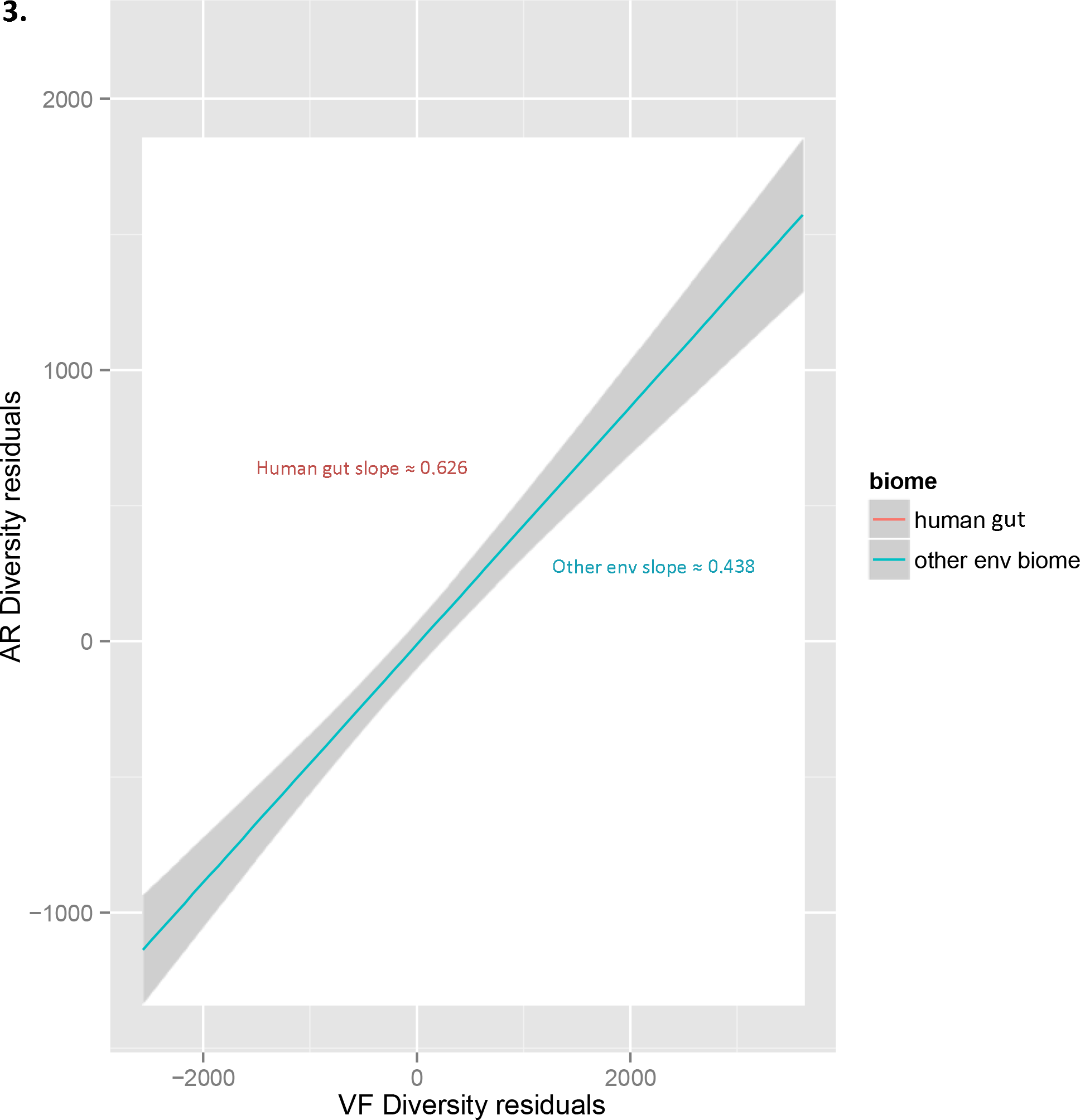
Distribution of AR by VF diversity in environmental metagenomes. Scatter plot ARd residuals versus VFd residuals of each environmental and human faeces 61 metagenomes (see Material and Method). Residuals are taken from the regressions shown in Figure 1.A and 1.B. Each dot represents a metagenome. The green line represents the regression line of ARd residuals environmental metagenomes on VFd residuals (the grey shading is the 95% confidence interval) and its slope is 0.44. The red line represents the regression line of ARd residuals of human faeces metagenomes on VFd residuals and the slope is 0.63. The difference in slopes is significant, but not very significant, with p-value of 3e-02.

Fig 4 shows residuals of AR diversity versus the residuals of VF diversity in the human gut metagenomes for each population (country) and for the three populations together. The partial Pearson correlation coefficient between ARd and VFd given metagenome richness is lower in the human gut metagenomes dataset (r=0.68, p-value 3e-22~0.0, Fig 4) than in the environmental samples (r=0.86 as seen above, Fig 3).

**Figure 4.**
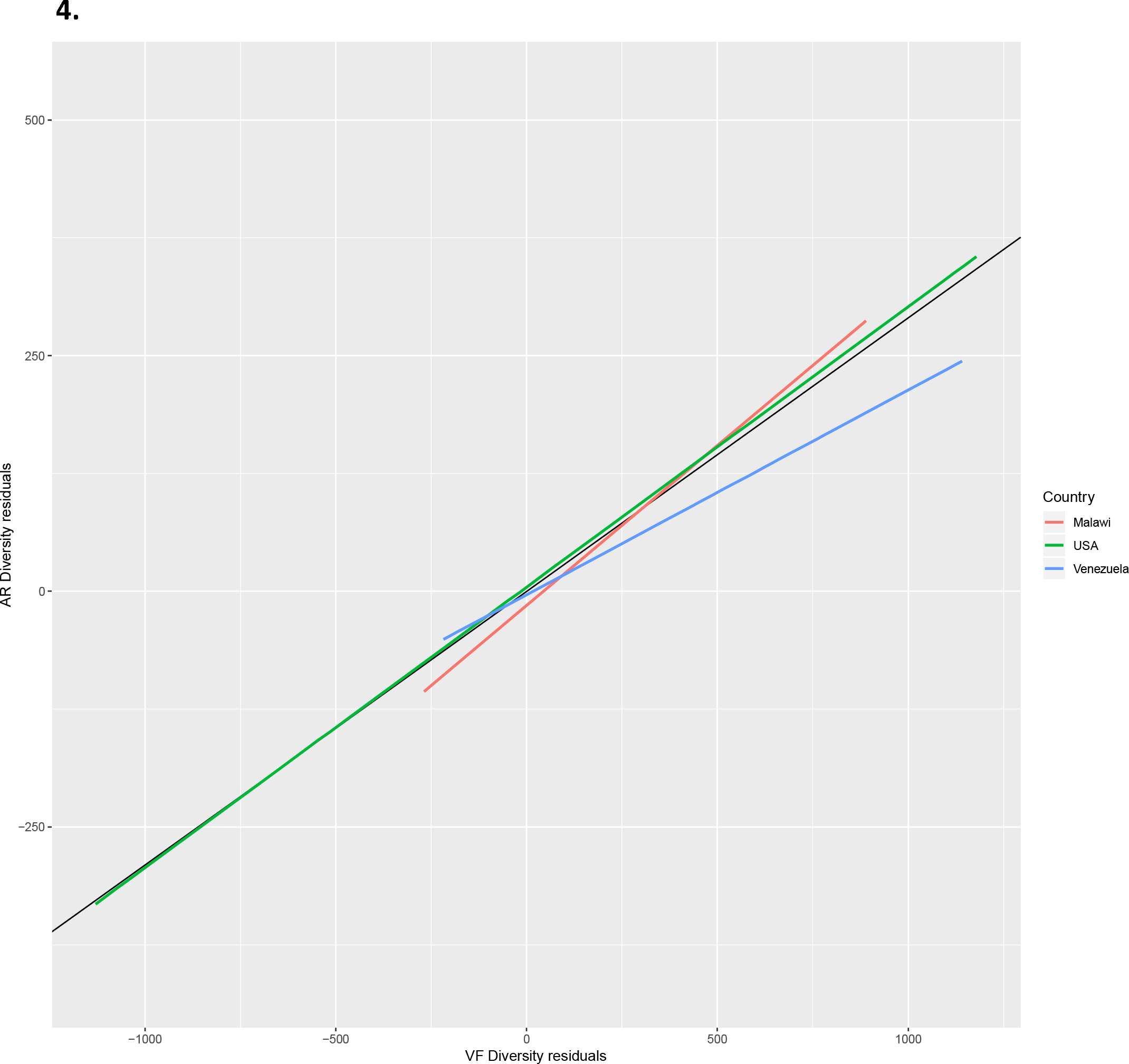
Distribution of AR by VF diversity in Human gut metagenomes. Scatter plot ARd residuals versus VFd residuals of each human gut 110 metagesnomes. Residuals are taken from the regressions shown in Figure 2.A and 2.B. Each dot represents a metagenome. In 2.A (resp.2B), the black lines represent the regression of the ARd residuals on VFd residuals for all the metagenomes. The green line represents the linear regression of the ARd residuals on VFd residuals for the USA metagenomes subset, the red line is the regression line for the Malawian metagenomes, and the blue line is the one for the Venezuelan metagenomes.

We can further distinguish different trends upon geographical localization of the human populations under study. The largest contribution to this graph comes from the North American samples, which account for 66/110 (60%) of the individuals (Fig 4). The Amerindians (Venezuela) metagenomes account for 21/110 metagenomes (19%) and the Malawians for 23/110 (21%). These two non-western human populations have very contrasting lifestyles and exposure to antibiotics: the uncontacted Amerindians, harbouring a resistome similar to that of pre-antibiotic era; and those of the people from Malawi with a high prevalence of infectious diseases and widespread use of unprescribed antibiotics (25– 27).

The Pearson partial correlation coefficients for metagenomes in the three populations are positive and in the same range (North americans: 0.66, Malawians: 0.81, Amerindians: 0.70). Regressions of the residuals of ARd on the residuals of VFd (Fig 4) shows a smaller slope for Amerindians (0.20) than for USA (0.31) and Malawians (0.33), but the differences are not significant: p-values spray from 8e-02 for the difference in Malawians-Amerindians slopes to 7.2e-01 for the difference in Malawians-North americans slopes.

### The co-occurrence of AR and VF belonging to the cell envelope

Our results suggest that co-occurrence of AR and VF might be taking place amidst bacterial communities. We wondered, however, which were the genetic traits that were more prone to this effect. We have then computed the partial correlations, given the metagenome richness, between every AR and VF protein families in the metagenomes under study to generate the correlogram presented in Fig 5. The vast majority of associations falls into the functional category of multi-drug efflux pumps (AR’s) associated with either secretion systems, as well with iron uptake and adhesion mechanisms (VF’s), respectively (Fig 5). Amongst the most representative associations between AR and VF traits are those belonging to the cell envelope and the general secretion mechanisms. On the other hand, the AR protein families for beta-lactamases, and general beta-lactam resistance mechanisms, bore no statistically significant correlations whatsoever.

**Figure 5.**
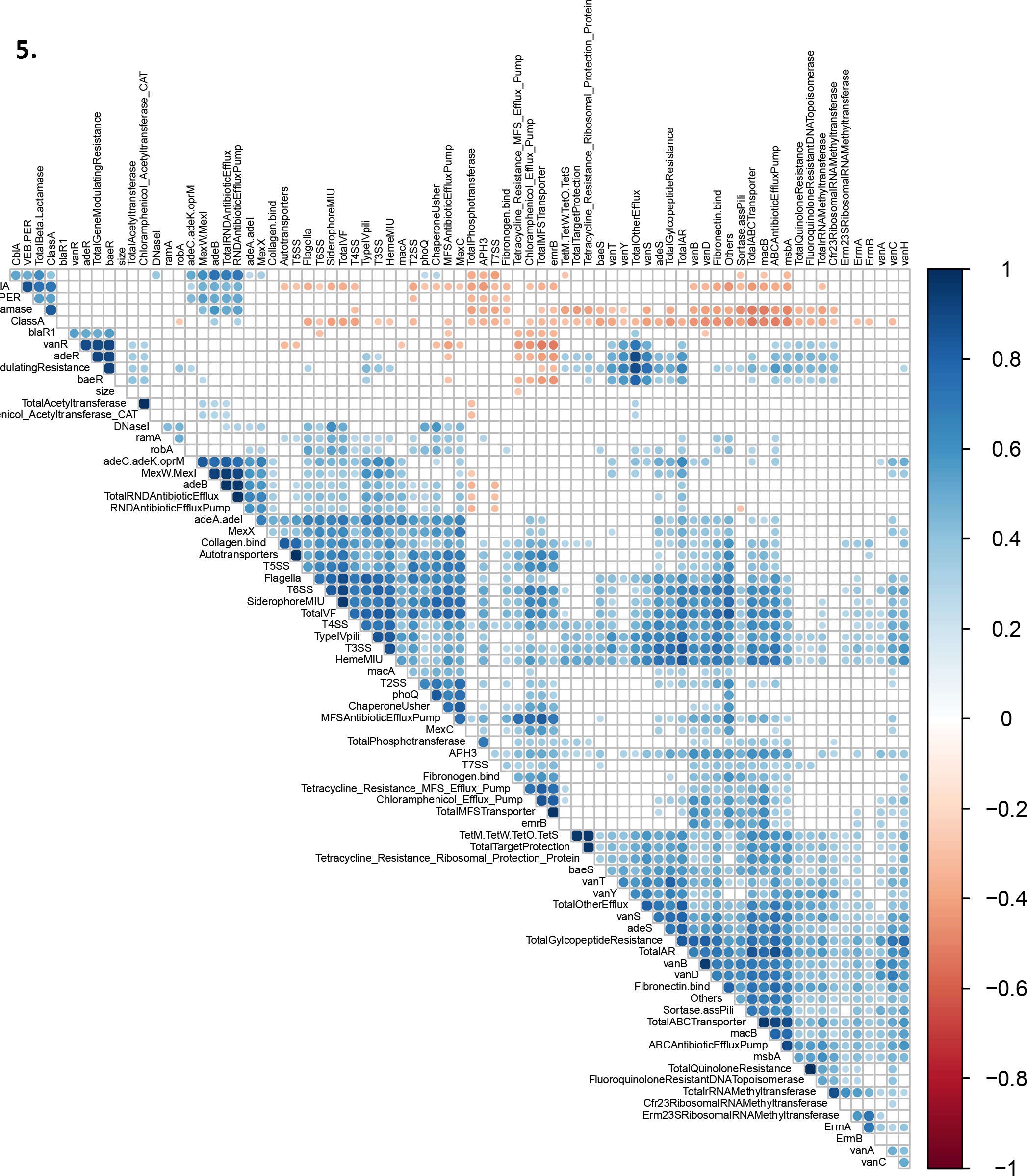
Correlogram of VR and AR protein families. Partial correlations for AR and VF protein families given metagenome richness have been computed. The color indicate negative (blue) or positive (red) partial correlation. Intensity of the color indicates the value. The point sizes indicate the significances of the partial correlation (white squares indicate no significant correlation).

## Discussion

### Antibiotic resistance among environmental and human gut metagenomes

As expected, we found different homologues for antibiotic resistance (AR) gene products belonging to different protein families (AR diversity or ARd) among environmental metagenomes (Fig 1A). We have shown that, in the environmental samples, ARd varies a lot from metagenome to metagenome. This may result from the differential microbial community composition of the metagenomes, whose genetic diversity can be grouped according to the adaptation to the environment in question (22); but also from the fact that the selective pressure for the maintenance of antibiotic resistance genes in environmental microbiome varies widely from environment to environment (28–30). Human gut metagenomes, on the contrary, have a less diversified repertoire of AR determinants (lower ARd) than that of their environmental counterparts.

We found a very strong correlation between the diversity of AR determinants and the metagenome’s protein family richness independently of the geographic origin of the human populations. These similar densities of ARd can indicate that, in human gut microbiomes, the number of different AR genes is not influenced by the human lifestyle, such as diet, medical care, access to antibiotics or other cultural habits (31), and/or that the adaptation to the intestinal tract shapes microbial AR diversity as well. This is a surprising results given that antibiotic use can vary from country to country (CDDEP), and, as consequence, individuals from different human populations are under different antibiotic exposition. Forslund and co-workers have demonstrated that there are robust differences in the antibiotic resistance arsenal between countries and that these differences follow the veterinary and human antibiotic use (27).

### Virulence among environmental and human gut metagenomes

In what concerns virulence, we can assert that there is a wide diversity and density differences of VFs in environmental metagenomes, which poses as evidence of the plasticity portrayed by environmental bacteria to adapt to different hosts and niches (Fig 1B). On the other hand, human gut microbiomes harbour a less diverse VF repertoire, especially in the US samples, which seem to indicate an evolution towards adaptation to the human gut, or lower contact with pathogens, eventually due to vaccinations and sanitation.

### Association of AR / VF

According to Yatsunenko and colleagues, once the human gut metagenome is established at the age of 3 years old, it does not diverge much from individual to individual, in what concerns both phylogenetic diversity and functional richness (23).

The most relevant results shown here are that, when corrected for metagenome’s protein family richness, ARd and VFd are strongly correlated both in the environmental samples (Figs 3) and in the human gut samples (Fig 4), with special emphasis on the human gut.

The North American intestinal samples (Fig 4) show a wide variety of associations between ARd and VFd, always presenting a statistically significant correlation between these genetic traits. This result, in itself, reinforces our hypothesis that antibiotic resistance and virulence are in fact co-associated, suggesting that the mobilization of mobile genetic elements as gene cassettes, driving to multirresistance, might be taking place. The US population, like other industrialized countries, is culturally exposed to antibiotics from health care facilities such as hospitals, antibiotic therapy, and the use of antibiotics in agriculture and livestock.

Malawians and amerindians from Venezuela share, phylogenetically, more similar human gut microbiomes than the North Americans (23), which belong mainly to a different enterotype (23, 32). Yet, whereas there is no ARd/VFd correlation among the Venezuelan individuals, there is a very strong correlation among the Malawian ones. This result highlights the potential effect of the exposure to antibacterial drugs on promoting the dissemination of antibiotic resistance by horizontal gene transfer, and the potential co-selection of virulence traits within bacterial communities. The uncontacted Amerindian microbiome represents a “frozen” relic of a pre-antibiotic era of the human resistome, while the Malawian gut microbiome is much more exposed, both to antibiotics, and to colonization by pathogens (25–27).

Amerindians have no known access to pharmaceutical drugs, as they usually make use of the traditional indigenous medicine. Malawi is one of the poorest countries in Africa, where most people live with less than one dollar a day, many people cope with AIDS and bacterial infections, and where many children suffer from severe malnutrition (25). Nutrition has been reported to have a big impact on both the human gut microbiome composition and resistome (31, 33, 34). In Malawi, UNICEF has been implementing a program of Ready-to-Use-Therapeutic-Food (RUTF) to reduce mortality among children. RUTF often contains antibiotics such as co-trimoxazole. It has been questioned, however, whether the success of this therapy is due to re-nutrition or to the combination with antibiotics (35, 36) that could have an effect on the microbial gut composition. It has been reported that there is widespread resistance to almost all of antibiotics that are empirically used in Malawi due to the lack of routine microbiological culture and sensitivity testing (36), but also due to self-medication. Human gut samples from the Malawi people of the dataset used in the present study are described as having a high overall resistance potential with an over representation of cephalosporin and tetracycline resistance genes which may suggest extensive use of old, broad-spectrum antibiotics, a known problem in many developing countries (27).

### The co-representation of AR and VF targeting the cell envelope

Between all the possible statistical AR and VF protein families’ correlations, the strongest ones are among proteins belonging to the bacterial cell envelope (Fig 5). This result is not surprising, as many proteins that belong to the secretory system or to the secretome itself, and thus target the bacterial envelope, are frequently encoded on mobile genetic elements (6). As such, this co-occurrence may also be regarded as a direct consequence of the dynamics of these genetic elements coding for both types of determinants.

In human gut, the strongest correlations involve efflux pumps that can extrude antibiotics non-specifically, and adhesion and iron scavenging mechanisms of pathogenicity. One possible explanation for the fact that the most frequent AR and VF associations in human gut involve efflux pumps could be that they allow a fast and efficient response to new man-made antibiotic molecules, while in environmental genomes, specific resistance mechanisms targeted to specific antibiotic molecules may have had more time to evolve to specific strategies.

Another possible explanation is that extrusion can also be implicated in pathogenic mechanisms of human associated pathogens as well as bacteria-host interactions. For example, biofilm formation in a *Staphylococcus aureus* methicillin resistant (MRSA) strain has been known to be essentially reliant on the activity of fibronectin binding proteins (37), and multi-drug efflux pumps have direct implications with the formation and maintenance of such biofilms (38). Furthermore quinolone resistant *S. aureus* strains up regulate the production of fibronectin binding proteins when subjected to sub-lethal dosages of ciprofloxacin (39). It has also been acknowledged that physiological levels of some cations present within the host promote the up regulation of genes encoding putative efflux transporters, oxidoreductases, and mechanisms of iron uptake either in *Acinetobacter baumannii* (40) as in *Burkholderia cenocepacia* (41), which could explain the co-association of iron acquisition systems with those of multi-drug efflux pumps.

We show here that, even after correcting for protein family richness, there is a correlation between the diversity of antibiotic resistance and virulence traits in human gut microbiomes and in environmental microbiomes. Interestingly, this correlation varies from a human population to another. Human antibiotic exposure due, either to therapy or to the environment and food, can have an effect on selecting for potentially pathogenic bacteria in the human gut microbiomes. It can also drive and shape changes on the gene pool of microbiomes. This means that, by selecting for resistant bacteria, we might be also selecting for more virulent strains, as a side effect of antimicrobial therapy.

## Materials and methods

### Metagenomic datasets

Our human gut query cohort included 110 previously studied, and publicly available metagenomes pertaining to individuals from different regions of Venezuela, Malawi, and the US, as well as a broad age span (0.05 to 53 years) (23) (dataset of the project mgp98 “Human gut microbiome viewed across age and geography (WGS)” at the MG-RAST metagenomic analysis server). All the metagenomes files were generated by the same team and project, and by using the same bioinformatics pipeline. Our environmental cohort comprised 64 previously selected, and publicly available environmental metagenomes, belonging to 12 different biomes (dataset collection of the metagenomes referred in the “Metagenomic mining for microbiologists” study) (22). Although Delmont’s team report using a dataset comprised of 77 metagenomes, there are only 70 MG-RAST accession numbers present in the article’s appendix, of which only 64 are publicly available. Both project’s metagenomes were downloaded from MG-RAST on the 29th of January 2016, under FASTA format, making use of successive calls to its Application Programming Interface (API) (42) using the respective MG-RAST’s accession numbers available in the aforementioned bibliography. Each FASTA file comprised clustered protein-coding sequences, retrieved from MG-RAST’s file-formatting pipeline (550.cluster.aa90.faa files). The protein sequences enclosed in these files are clustered at 90% identity, containing non-redundant translated sequences. These protein-coding formatted FASTA files contain the translation of one representative from each cluster. Thus, the numbers of protein sequences for a given metagenome used here represent its richness.

### BLAST, VFDB and Resfams

For every metagenome present in our query, a BLASTP (43) search was performed against the VFDB database for bacterial virulence factors families (Virulence Factors of Pathogenic Bacteria VF.pfasta dataset of the Institute of Pathogen Biology, China) (24) and the Resfams AR Proteins database of bacterial antibiotic resistance protein families (Resfams AR Proteins - v1.2 database of the Dantas Lab at the Washington University School of Medicine, St Louis) (44). An alternative approach made use of the Antibiotic Resistance Genes Database (ARDB) (reference antibiotic resistance genes dataset resisGenes.pfasta of the Center for Bioinformatics and Computational Biology University of Maryland) (45), but several hindrances concerning its sub-classification by functional antibiotic resistance protein families made us discard the possibility of using such database to address of AR and VF protein family co-occurrence in metagenomes.

The BLAST+ executables package was downloaded on the 17th of November 2015 (ncbi-blast-2.2.31+ version). The VFDB was downloaded on the 11th of November 2013 (31 classified FASTA files of bacterial virulence factor sub-families), and the Resfams database was downloaded on the 29th of January 2016, (123 classified FASTA files of bacterial antibiotic resistance protein subfamilies. Every protein enclosed in each of the addressed metagenomes, was used as a query in order to search for similarities to either AR or VF protein-coding traits. Hence, we aimed at retrieving the best hit (best scored alignment) that enabled us to assign an AR or VF function to each of the aforementioned metagenomic proteins. Every BLASTP search was performed with a very stringent e-value cut-off of 1e-15 (10 orders of magnitude lower than the conservative commonly used E-value of 1e-5). Next, we filtered the resulting output files, as to only retrieve the alignment hits with >60% coverage of the query size, and whose query and subject length ratio is between 0.75 and 1.5. When using a combination of this very stringent e-value of 1e-15 together with coverage, and length ratio filters, we expect to only retrieve true homologues, avoiding false positives. Furthermore, from all the generated alignments between our metagenomic query cohort and the preceding databases, we computed the representative counts for the different gene families that coded for either AR or VF traits. Thus, the amounts of different classes (gene families) that are present in a given metagenome represent its diversity in terms of AR or VF traits.

We have also removed hits for proteins that aligned with both antibiotic resistance and virulence factor proteins (25.9% of hits against the VFDB, and 29,9% of the hits against the Resfams database). The abovementioned filters and algorithms were implemented making use of Unix scripting languages (GNU Awk version 4.0.1, and Z-Shell version 5.0.2), under a Linux environment.

### Statistical Analysis

To test for relationships in metagenomes between the presence of both antibiotic resistance and virulence factors traits, we proceeded as follows. Our expectation is that the diversity number for homologues of antibiotic resistance protein families (henceforth denoted as ARd) and the diversity number for homologues of virulence factors protein families (denoted as VFd) in each metagenome (clustered at 90% identity) increases with the protein family richness of the latter (being that the protein family richness here means the total number of cluster representative proteins). Given the potential diversity of these protein families, one does not expect ARd and VFd to level off with a metagenome’s protein family richness. Therefore, we assume a linear relationship between ARd and metagenome’s protein family richness, with a fixed 0 intercept. The same for VFd. Our results will show that these assumptions are reasonable. Thus, to avoid spurious correlation, we corrected the diversity of ARd and VFd in a metagenome by taking into account its protein family richness: we have computed the partial correlations between ARd and VFd given the metagenome richness, and equivalently, when plotting ARd versus VFd, we have instead taken residuals of regression of ARd on metagenome richness and residuals of regression of VFd on metagenome richness. Recall that the partial correlation measures the degree of association between two random variables, with the effect of a set of controlling random variables removed. For example, the partial correlation between A and B given a third variable C correlated with A and B, is the correlation between the residuals *e*_*a*_ and *e*_*b*_ resulting from the linear regression of A with C and of B with C. If r_AB_, r_AC_ and r_BC_ are the correlation coefficients between A and B, A and C, and B and C, the partial correlation between A and B giveN C, r_AB.C’_ can be computed as 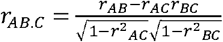.

## Acknowledge

We thank Eduardo P. C. Rocha for critical reading of the manuscript and constant interest and suggestions since the beginning of this work, Guillaume Sapriel for suggestions to improve the methods, Pedro David, Eric Duchaud, Guillaume Achaz, and all the members of our teams for the fruitful discussions, Octávio Paulo, Francisco Pina-Martins and Sophie Brouillet for the technical help, and granting access to the servers.

Conceived and designed the experiments: JP TN. Performed the experiments: PE. Analysed the data: JP FD TN. Wrote the paper: JP FD TN.

